# Assessing a Role for Sarcospan in Immune Function

**DOI:** 10.1101/2024.03.31.587501

**Authors:** Isela C. Valera, Aida Rahimi Kahmini, Kyle A. Smith, Rhiannon Q. Crawford, Michelle S. Parvatiyar

**Author notes:** Correspondence should be addressed to: Michelle S. Parvatiyar, Florida State University, Department of Health, Nutrition, and Food Sciences, Biomedical Research Facility, 107 Chieftan Way, 238, Tallahassee, FL 32312.

## Abstract

Sarcospan (SSPN) is a tetraspanin-like member of the dystrophin-glycoprotein complex (DGC) with expression reported at the transcript level in B cells and several other immune cell populations. SSPN is best known for its expression in skeletal, cardiac, and vascular smooth muscle, however it is expressed in other cell types and its roles in non-muscle cells have not been well described. Analysis of functional interactions of SSPN with effectors of hematopoietic cell lineages uncovered strong associations with several proteins with known roles in hematopoiesis and immune response. In this study, we utilized flow cytometry to uncover SSPN+ immune cell populations expressing the SSPN protein. We performed immunophenotyping of global SSPN-deficient (SSPN^-/-^) C57BL/6J male and female mice to assess whether SSPN plays an essential role in generation of specific immune cells. By flow cytometry we found that B cells, CD11b+ and CD11c+ (monocytes, tissue macrophages, and a subset of dendritic cells) express the highest levels of SSPN at their cell membranes. To pinpoint potential roles for SSPN in B cell development, SSPN^-/-^ bone marrow cells were isolated and differentiated using phorbol myristate acetate (PMA). To test B cell effector functions, antibody isotype analysis was performed to assess whether SSPN influences antibody class switching. Blood serum of WT and SSPN^-/-^ mice were tested for presence of important developmental signals for B cell maturation, development and isotype switching. Our study is the first to document SSPN protein expression on distinct classes of immune cells including CD11b+, CD11c+ and B cells (CD19^+^). Immunophenotyping revealed several differences in immune cell populations in the murine spleens: 1) a significant increase in % T cells in SSPN^-/-^ males, 2) slight reduction of % B cells in SSPN^-/-^ females, 3) a significant increase % of dendritic cells in SSPN^-/-^ females, and 4) no significant alterations in NK cells. Under unstimulated conditions, SSPN^-/-^ female mice had a significantly lower IgG2b titers compared to WT. Overall, the observation of SSPN expression in distinct immune cells, potential role(s) in hematopoiesis and B cell effector function is intriguing and warrants further investigation.

**Highlights:** The sarcospan protein has an established role in stabilization of striated muscle membranes and is an essential member of the dystrophin-glycoprotein complex. This study investigates the reported presence of *Sspn* in B cells and examines potential roles in immune function including:

1. Determining a connection with hematopoiesis and establishment of ‘niches’
2. Impact on immune content in spleens of sarcospan-deficient mice
3. Identifying a role in B cell effector functions

## 1. Introduction

Sarcospan (SSPN) is a tetraspanin-like protein that has been found distributed in different tissues throughout the body. In the heart, skeletal muscle, and vascular tissue it has been shown to exist within the dystrophin-glycoprotein complex (DGC) where it helps to maintain integrity of the complex and sarcolemma stability [1-5]. Mutations in dystrophin cause loss of the protein that can manifest as Duchenne muscular dystrophy (DMD), which causes membrane instability that leads to muscle pathology, muscle weakness and activation of immunological and inflammatory processes [6]. The role of SSPN in other cell types is less known although the presence of *Sspn* RNA has been found in other tissues [1] with a tissue distribution that closely matches dystroglycan [7]. *Sspn* RNA expression has been observed in immune cells with the highest expression in B cells (The Human Protein Atlas). There is limited data concerning involvement of SSPN in the complex signaling that underlies normal immune cell development and immune response. SSPN has structural similarity to tetraspanin proteins, which have been shown to have an important role in immune cell recruitment and migration. Tetraspanins have been shown to organize binding partners within cell membranes and enable immune cell migration throughout the body [8].

The immune system is comprised of two major branches, the innate and adaptive systems, each contributing their own contrasting separate arms of the host response. Significant synergy exists between the two systems whereas the innate immune system contributes to activation of the antigen-specific cells within the adaptive system, whereas adaptive responses are amplified by recruitment of innate effector mechanisms. The innate immune system is comprised of all the host immune defense mechanisms encoded by germ-line genes, is fast-acting (within hours), and is considered the first line of host defense. Cells of the innate immune system typically arise from a common myeloid progenitor that is derived from a multipotential hematopoietic stem cell in the red bone marrow. Innate immune cells include dendritic cells, macrophages, neutrophils, basophils, eosinophils, and mast cells that produce cytokines or interact with other cells ultimately activating the adaptive immune system. In addition, natural killer (NK) cells are innate lymphoid cells that act as cytotoxic effectors and regulators of the immune response.

The adaptive immune system, however, exhibits a slower response (several days) after its effector cells undergo specificity for its target antigens. Antigens are immunogens that trigger the immune system. Adaptive immune responses depend on T and B lymphocytes which express antigen-specific receptors that arise from the same common lymphoid progenitor as NK cells. Antigen-specific receptors are encoded by genes that undergo somatic rearrangement of germ-line gene elements to form highly specific intact T cell receptor (TCR) and immunoglobulin B cell antigen receptor, Ig genes. The ability to assemble a vast number of antigen receptors from a limited number of germ-line-encoded gene elements allows the adaptive system to mount a highly specific response to foreign antigens. B cells play an important role in mediating humoral immunity that involves generation of antibody producing plasma cells and memory B cells. Transitional B cells provide a link between immature and mature B cells and travel from the bone marrow to secondary lymphoid tissues [9]. *Sspn* expression is very low in naïve B cells and increased in activated B cells.

This is the first study examining SSPN protein expression in immune cells and the immune cell repertoire in the absence of SSPN. This study was initiated to investigate and potentially understand the functional consequences of *Sspn* transcripts in B cells and other immune cells to determine whether they express SSPN protein. We also investigated a potential role for SSPN in several important immune functions including B cell proliferation or immunoglobulin class switching. Immunophenotyping was performed to determine whether deletion of the *SSPN* gene alters immune signaling and subsequently immune cell populations in SSPN^-/-^ mice.

## 2. Methods

### 2.1. Mouse Models

We utilized WT (C57BL6/J) (JAX#000664) and Sspn knock-out mice (JAX#006837) obtained from Jackson laboratories and generated by the Campbell laboratory [10]. In our colony SSPN-deficient (SSPN^-/-^) mice were backcrossed twice on C57BL/6J mice and progeny obtained were used to maintain male and female WT and SSPN^-/-^ mice used for this study. Mice were utilized at 3-4 months of age and fed standard chow diet LabDiet #5001-RHI-E 14 (4.5% crude fat) ad libitum.

### 2.2. Flow Cytometry and Sample Preparation

#### Sample Preparation

Spleens were collected from WT and SSPN^-/-^ mice and weighed. Mouse body weight and tibia lengths were also recorded. Single cell suspensions were incubated for 30 minutes on a rocker at room temperature (RT) then filtered through a 70 μm strainer, washed and centrifuged. ACK lysis buffer was added to remove red blood cells and splenocytes were incubated with antibodies against CD16/32 for Fc receptors blockage for at least 30 min prior to staining. After blocking, the cells were incubated with a combination of fluorescently labeled anti-mouse antibodies.

#### Antibody Labeling

Cells were incubated with anti-mouse primary antibodies to assess immune populations with the following conjugated antibodies: B220-APC (Tonbo Biosciences, San Diego, CA; clone RA3-6B2), CD19-PE/Dazzle594 (BioLegend, San Diego, CA; clone 6D5), CD3-AlexaFluor488 (BioLegend, clone 17A2), CD4-violetFluor450 (Tonbo Biosciences, clone GK1.5), CD8-PE/Cy7 (Tonbo Biosciences, clone 53-6.7), NK1.1-PE (Tonbo Biosciences, clone PK136), CD335/NKp46-APC (BioLegend, clone 29A1.4), CD11b-APC/Cy7 (Tonbo Biosciences, clone M1/70), CD11c-PerCp/Cy5.5 (Tonbo Biosciences, clone N418), Ly6G-PE (Tonbo Biosciences, clone RB6-8C5), and F4/80-FITC (Tonbo Biosciences, clone BM8.1,). Relevant isotype controls were used for each experiment. SSPN was measured using the primary unconjugated anti-SSPN antibody (Santa Cruz, sc-393187).

#### Analysis

Samples were run on either FACSCalibur or FACSCanto flow cytometers (BD Biosciences) and spectra analyzed using either FlowJo (Becton, Dickinson & Co, Ashland, OR) Dor FlowLogic software (Inivai Technologies, Mentone, Australia).

### 2.3. Tissue Histology

Spleens were isolated, embedded in OCT freezing medium, and flash frozen in N_2_ cooled isopentane. The frozen tissue was sectioned into 7 μm thick sections and placed onto charged microscope slides. The sections were then stained using H&E to visualize the histological features of WT and SSPN^-/-^ spleens. Brightfield images were obtained using a Leica DMi8 microscope at 20x magnification.

### 2.4. Proliferation Assay

Mouse spleens were harvested, and single cell suspension prepared in RPMI complete medium at approximately 5-10 x 10^6^ cells/ml. Proliferation of mouse splenocytes was stimulated by addition of 10 ng/ml phorbol 12-myristate 13-acetate (PMA) directly to the cell suspensions to activate both myeloid and lymphoid lineages and total cells counted. Supernatants were collected for cytokine analysis. Flow cytometry was performed to assess the amount of proliferation in WT and SSPN^-/-^ cultures in response to PMA was determined using carboxifluorescein diacetate succinimidyl ester (CFSE) dilution and anti-CD19 antibodies to determine the signal detected by isotype control -PMA, isotype control + PMA, antibody - PMA, and antibody + PMA.

### 2.5. Immunoglobulin Isotype Analysis

Blood serum was collected from WT and SSPN^-/-^ mice and allowed to clot for at least 30 mins prior to centrifugation for 10 mins at 1,000 x g. Serum was aliquoted and stored at -20ºC until use. Thawed samples were centrifuged to remove debris. The antibody-immobilized beads were resuspended using sonication. The LEGENDplex™ Mouse Immunoglobulin Isotyping Panel catalog #740493 with the V-bottom plate was used according to manufacturer’s instructions (BioLegend, CA). Briefly, standards were prepared fresh by reconstitution of lyophilized mouse immunoglobulin isotyping panel standard cocktail. Samples and standards were run in duplicate, incubated for 2 hrs RT with mixed beads with agitation and protected from light. Supernatant was removed and beads were washed twice prior to addition of Detection Antibodies and incubated at RT for 1 hr. Secondary antibody SA-PE was added to each well incubated at RT for 30 min, samples centrifuged and washed, and beads resuspended and read by flow cytometry.

### 2.6. Analysis of B cell Functions

To assess whether loss of the SSPN protein affects essential immune cell functions spleens were harvested and a single cell splenocyte suspension was prepared in RPMI complete medium (10% FBS (GenClone 25-550H), 1% pen/strep (GenClone25-512), 50 μM 2-ME (Sigma 60-24-2). Cell Stimulation Cocktail (Tonbo # TNB-4975-UL100) was directly added (1X) (2 μL/mL) to the cell suspension. The splenocytes were cultured overnight (12 hours) at 37°C, 5% CO_2_ and cell media collected for analysis. Levels of cytokines in the media were assessed using the LEGENDplex Mouse B cell multiplex bead-based panel w/VbP according to manufacturer’s instructions. This included B cell activating factor (BAFF), B cell maturation antigen (BMCA), interleukin 6 (IL-6), and transforming growth factor-beta (TGF-β) levels, which have important roles for B cell function, activation, and survival.

#### Statistics

All values in the text and figures are presented as mean +/- SEM, unless otherwise indicated. Statistical significance was determined using a two-tailed Student’s t-test to compare two relevant groups. Ordinary one-way ANOVA was used when comparisons were made across all groups and several sex differences were noted, followed by Tukey’s multiple comparisons test. *p* values of ≤ 0.05 were considered significant.

#### Study Approval

The animal studies included in this study were reviewed and approved by the Institutional Animal Care and Use Committee at Florida State University as protocol #IPROTO202200000003.

## 3. Results

### 3.1. Assessment of Functional Interactions between SSPN and Networks Involved in Hematopoietic Cell Lineages

To assess a role for SSPN in immune-related functional networks the ImmuNet web interface was examined to find networks SSPN had strong functional relationships [11]. The ImmuNet hematopoietic cell lineage data base provides information concerning direct and indirect functional relationships (Figure 1A). Seven of these have direct functional interactions with SSPN including CTSO (cathepsin O), RBMS3 (RNA binding motif, single stranded interacting protein 3), ARHGEF6 (rac/cdc42 guanine nucleotide exchange factor (GEF) 6), ID4 (inhibitor of DNA binding 4, dominant negative helix-loop-helix protein), FBN1 (fibrillin 1), LRRC17 (leucine rich repeat containing protein), MFAP5 (microfibrillar associated protein 5). Eleven functionally interacting proteins with SSPN are shown with higher than 0.89 minimum relationship confidence as shown in Figure 1B. Each of these may provide insight into diverse roles of SSPN including response to cancer drugs, immune response to infection, blood disorders and susceptibility to lymphomas and other cancers.

**Figure 1.**
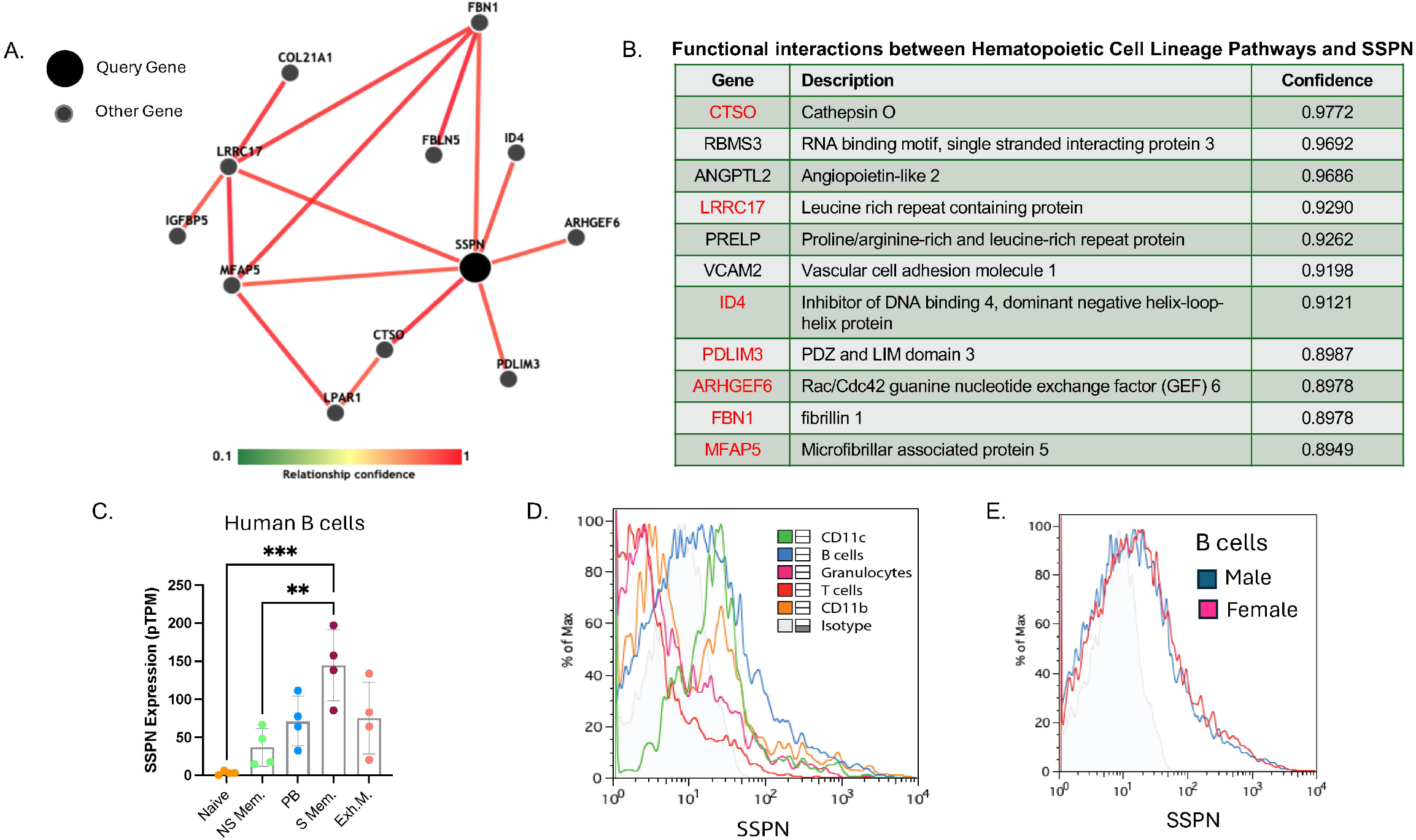
Evaluating Roles for Sarcospan in Immune Cell Function and Hematopoietic Cell Lineages. (A) Immune-related functional relationship networks were examined using ImmuNet and shown for the SSPN protein in the hematopoietic cell lineage dataset at a confidence level > 0.8949. (B) The table shows the eleven strongest functional relationships between hematopoietic cell lineages and SSPN. In (A) extracellular flow cytometry was utilized to identify SSPN-expressing immune cells in WT C57BL6/J mice. SSPN was detected on unstimulated B cells, CD11b+, and CD11c+ cells while unstimulated granulocytes and T cells had either no or low SSPN expression. In (B) B cells obtained from WT male and female mice exhibit similar SSPN expression profiles. In (C) RNA-seq data (nTPM) is plotted from the Human Protein Atlas for different B cell types (naïve, non-switched memory, switched memory, plasmablast, exhausted memory) (Monaco et al. 2019).

### 3.2. Analysis of SSPN Protein Expression in Immune Cell Types

To determine whether the SSPN protein is present in the plasma membranes of immune cells we utilized flow cytometry to detect SSPN expression in WT C57BL6/J splenocytes. SSPN expressing immune cells were identified by the presence of extracellular markers including CD11c for dendritic cells, CD19 to identify B cells, Gr-1 antigen to identify granulocytes, CD4 and CD8 to identify T cells, CD11b a leukocyte-specific marker for identification of monocytes/macrophages and natural killer cells. Flow cytometry results using male WT C57BL6/J splenocytes are shown in Figure 1C, and several cell populations exhibited a rightward shift compared to the isotype control peak indicating SSPN expression. SSPN expressing immune cells observed included SSPN+CD19+, SSPN+CD11c+, and SSPN+CD11b+ cells. In Figure 1D it is shown that SSPN+CD19+ B cell populations in WT C57BL6/J splenocytes are similar between male and female mice.

In Figure 1B data is included from the Human Protein Atlas https://www.proteinatlas.org/ENSG00000123096-SSPN/immune+cell that shows *SSPN* gene expression in different B cell populations including naïve B cells [12,13], non-switched, switched and exhausted memory B cells, and plasmablasts [13]. Low levels of *SSPN* gene expression were also shown for eosinophils, neutrophils, progenitor cells, and total peripheral blood mononuclear cells (PBMC) [13] and basophils [12]. Original RNA-Seq data can be found in the report by Monaco *et al*. [13] and the DICE (database of immune cell expression) database [12].

### 3.3. Evaluation of Spleen Weights and Tissue Histology

Immune cell content was assessed in the spleens of WT and SSPN^-/-^ mice. First, spleen weights were assessed to determine whether any differences existed between male and female WT and SSPN^-/-^ mice. Overall, in Figure 2A no differences were noted in the spleen weight/tibia length values between genotype, however significant differences were found between males and females that is likely related to the estrus cycle. In Figure 2B H&E-stained spleen transverse sections are shown for WT and SSPN^-/-^ male mice. Overall, no obvious morphological differences were observed, although the area of darker stained white pulp, which contains lymphoid tissues, were visibly larger in the SSPN^-/-^ spleens. Discrete regions within the splenic follicles could not be well visualized, but these consist of the marginal zone containing B cells, macrophages, and dendritic cells. Germinal centers contain activated B cells and T cells are in a non-central location within the periarteriolar sheath.

**Figure 2.**
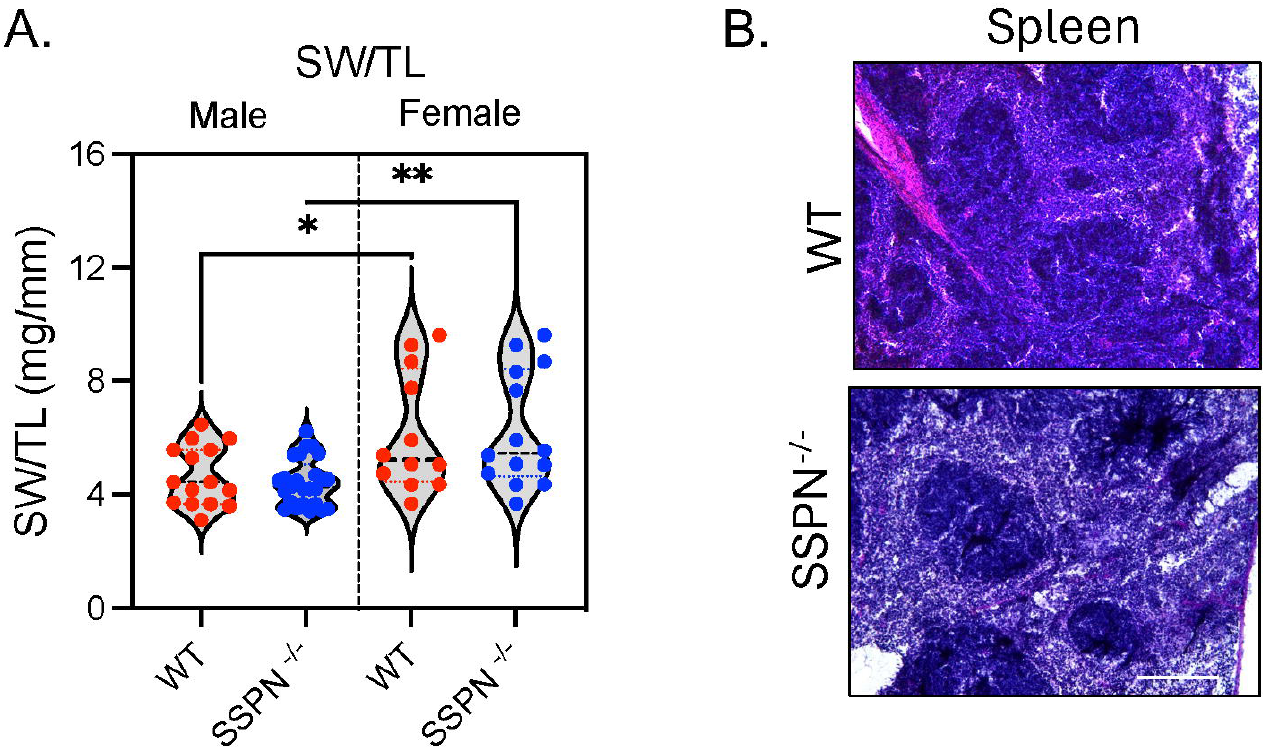
Determining the Impact of Sarcospan Loss on Spleen Morphology. In (A) spleen weight (grams)/tibia length (mm) (SWTL) are compared between WT and SSPN^-/-^ male and female mouse groups. (In (B) hematoxylin & eosin (H&E) stained transverse spleen tissue sections, images (20x). Bar = 50 μm. Data shown as individual values and statistics determined using one-way ANOVA followed by Tukey’s multiple comparisons test and considered significant if *p* ≥ 0.05. Significant differences indicated by * for 0.05 and ** for 0.001.

### 3.4. Assessment of B Cell Proliferation in SSPN-Deficient Splenocytes

Flow cytometry was used to assess whether SSPN deletion influenced B cell maturation altering the % CD19+B220+ B cells in SSPN^-/-^ versus WT splenocytes (Figure 3A and 3B). CD19 is expressed throughout B cell development (pro-B cells) and maintained throughout B cell differentiation up to the plasmablast stage (plasma cell terminal differentiation), while B220 is a pan B cell marker. Therefore, it appears that B cell development was not affected by the loss of SSPN, although female SSPN^-/-^ trended towards lower B cell counts, see Figure 3B.

**Figure 3.**
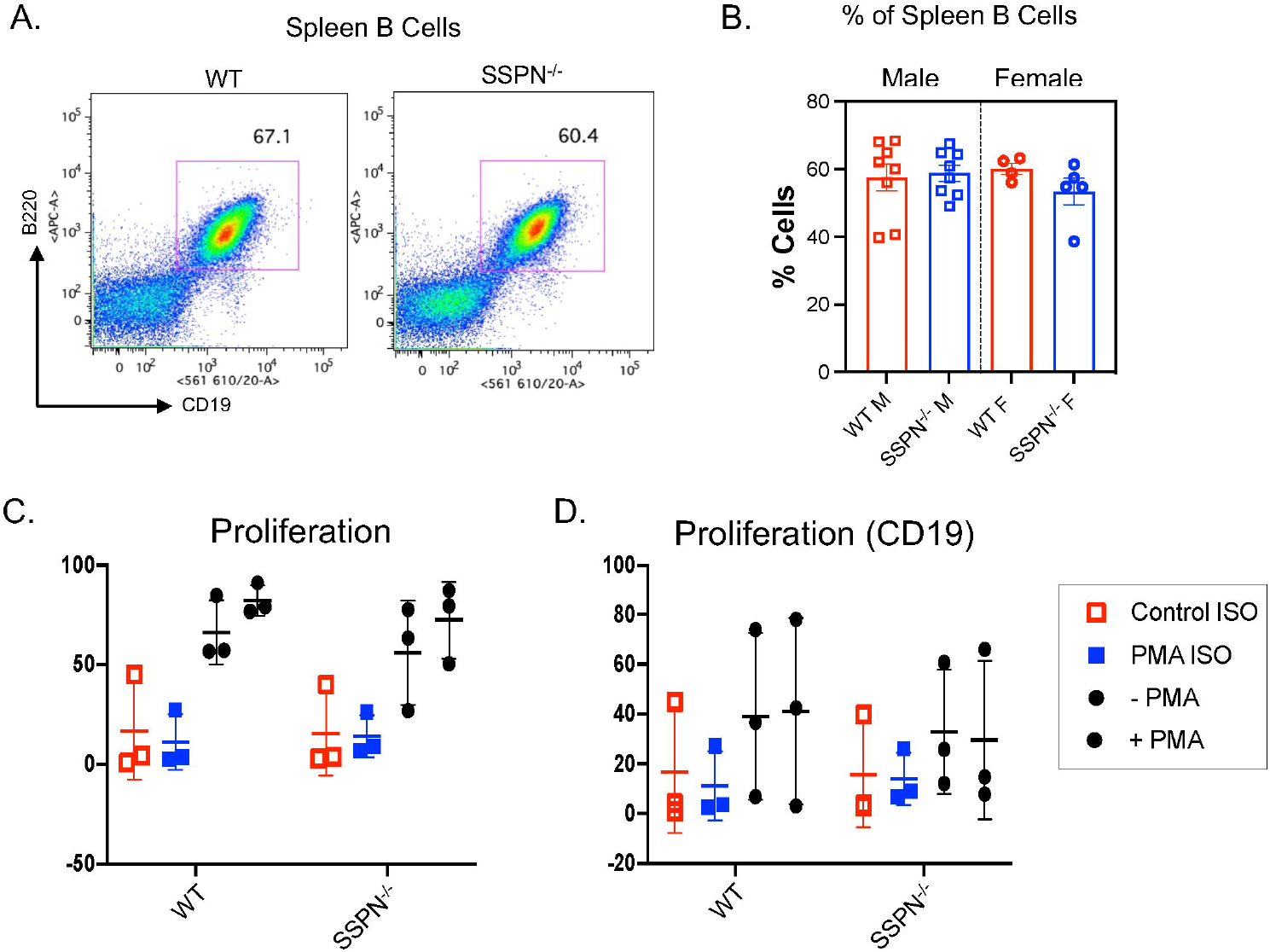
Examination of B Cell Proliferation in the Absence of Sarcospan. In (A) representative flow cytometry spectra of % B cells in WT and SSPN^-/-^ male mice. B) Representative flow spectrum shows B cell gating (B220^+^CD19^+^) in isolated splenocytes. (C) Shows % B cells found in male and female mice of both genotypes. A proliferation assay was conducted using splenocytes obtained from WT and SSPN^-/-^ mice using phorbol 12-myristate 13-acetate to stimulate cell proliferation of (C) all immune cells (pan immune cell marker) and (D) CD19^+^ B cells. Symbols indicate isotype control -PMA (red square), isotype control +PMA (blue square), antibody -PMA (black circle), and antibody +PMA (black circle). Data shown as individual values and statistics determined using one-way ANOVA followed by Tukey’s multiple comparisons test and considered significant if *p* ≥ 0.05. Significant differences indicated by * for 0.05 and ** for 0.001.

Splenocyte proliferation was assessed by flow cytometry to determine whether SSPN deletion affects overall proliferation using a pan immune cell marker as shown in Figure 3C and CD19+ B cell proliferation as shown in Figure 3D. Overall proliferation was evaluated + or – PMA and overall SSPN loss did significantly impact our findings. For the CD19+ cell population, there were no significant differences between + or – PMA groups of either genotype. There was a trend toward reduced CD19+ cell proliferation in splenocytes isolated from SSPN^-/-^ mice. However, there was no significant difference in immune cell proliferation and specifically B cell proliferation appeared not to be significantly affected by SSPN loss.

### 3.5. Immunophenotype Analysis of SSPN-Deficient Splenocytes

Immunophenotyping was performed to assess the impact of SSPN deletion on major immune cell populations. To assess this, splenocytes were isolated from WT and SSPN^-/-^ female and male mice. In Figure 4A the % of total CD4+ and CD8+ T cells is plotted, and the overall % of T cells was increased in male SSPN-/- splenocytes compared to WT. Figure 4D the representative flow images are shown indicating the populations of CD4+ and CD8+ T cells observed in WT and SSPN^-/-^ spleens. In Figure 4B the % of splenic plasmacytoid dendritic cells (pDC) were identified as B220+CD11c+ cells and representative flow spectrum shown in Figure 4E. Overall, there was a trend toward reduced numbers of these cell types in the SSPN^-/-^ male and female spleens compared to WT. In contrast, female SSPN^-/-^ mice exhibited a significant increase in % splenic dendritic cells (DCs) identified as CD11b+CD11c+ cells as shown in Figure 4C and representative flow images are shown in Figure 4F. Figure 4G shows representative images of flow spectra for NK1.1+ and NKp46+ cells and in Figure 3H the % of each cell population. Overall, there were no significant differences in either NK cell population.

**Figure 4.**
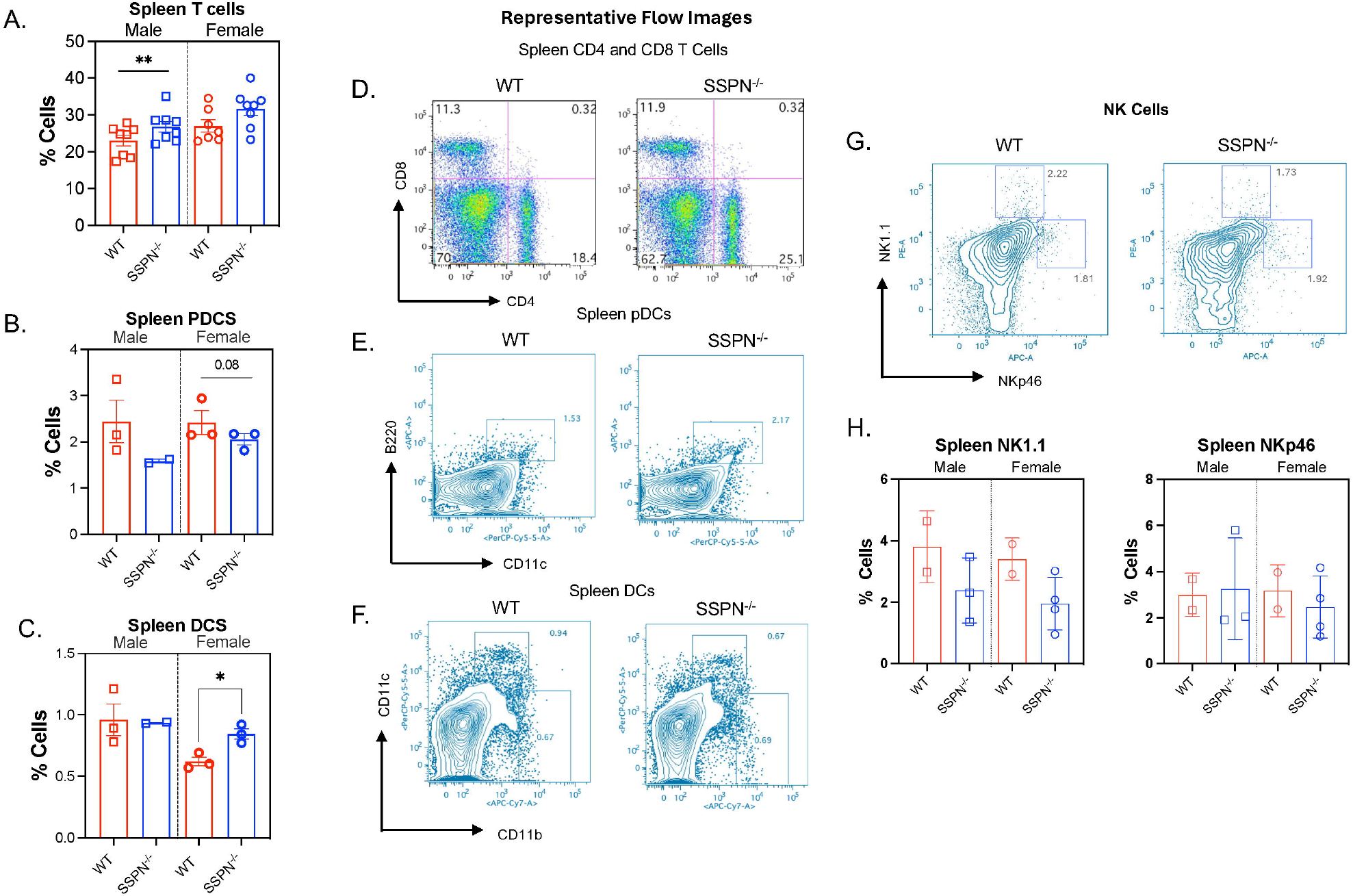
Immunophenotype Results for Male and Female Sarcospan-Deficient Mice. Flow cytometry was used to determine the % immune cell populations in splenocyte suspensions obtained from WT and SSPN^-/-^ male and female mice. In (A) T cells, (B) plasmacytoid dendritic cells (pDCs), (C) dendritic cells (DCs), and (G) natural killer NK1.1^+^ cells. Representative flow spectra are shown in (D) CD4+ and CD8+ T cells (E) pDCs, and (F) DCs. Genotype differences in the male and female mouse groups were assessed separately using the two-tailed Student’s t-test. Data shown as individual values and considered significant if *p* ≥ 0.05. Significant differences indicated by * for 0.05 and ** for 0.001.

### 3.6. Analysis of B cell Function and Survival in SSPN-Deficient Mice

To assess whether loss of SSPN protein affect aspects of B cell function, activation, and survival we examined class switching, an important B cell function, to determine whether it was affected by SSPN deletion. Immunoglobulin subtypes in the blood serum were compared between WT and SSPN^-/-^ mice in Figure 5A for males and Figure 5B for females. At steady state the most abundant isotypes in the blood serum of the mice were IgG1and IgG2b. In SSPN^-/-^ female mice there was a significant reduction in IgG2b isotype antibodies and an overall trend of lower IgM, IgG3, IgA isotypes compared to WT shown in Figure 5B. This is consistent with the reduction of B cell % in the SSPN^-/-^ female mice. For the SSPN^-/-^ female mouse there was one data set that was markedly different at baseline and at the edge of the plate. It was identified as an outlier by GraphPad as ROUT (Q = 0.5%).

**Figure 5.**
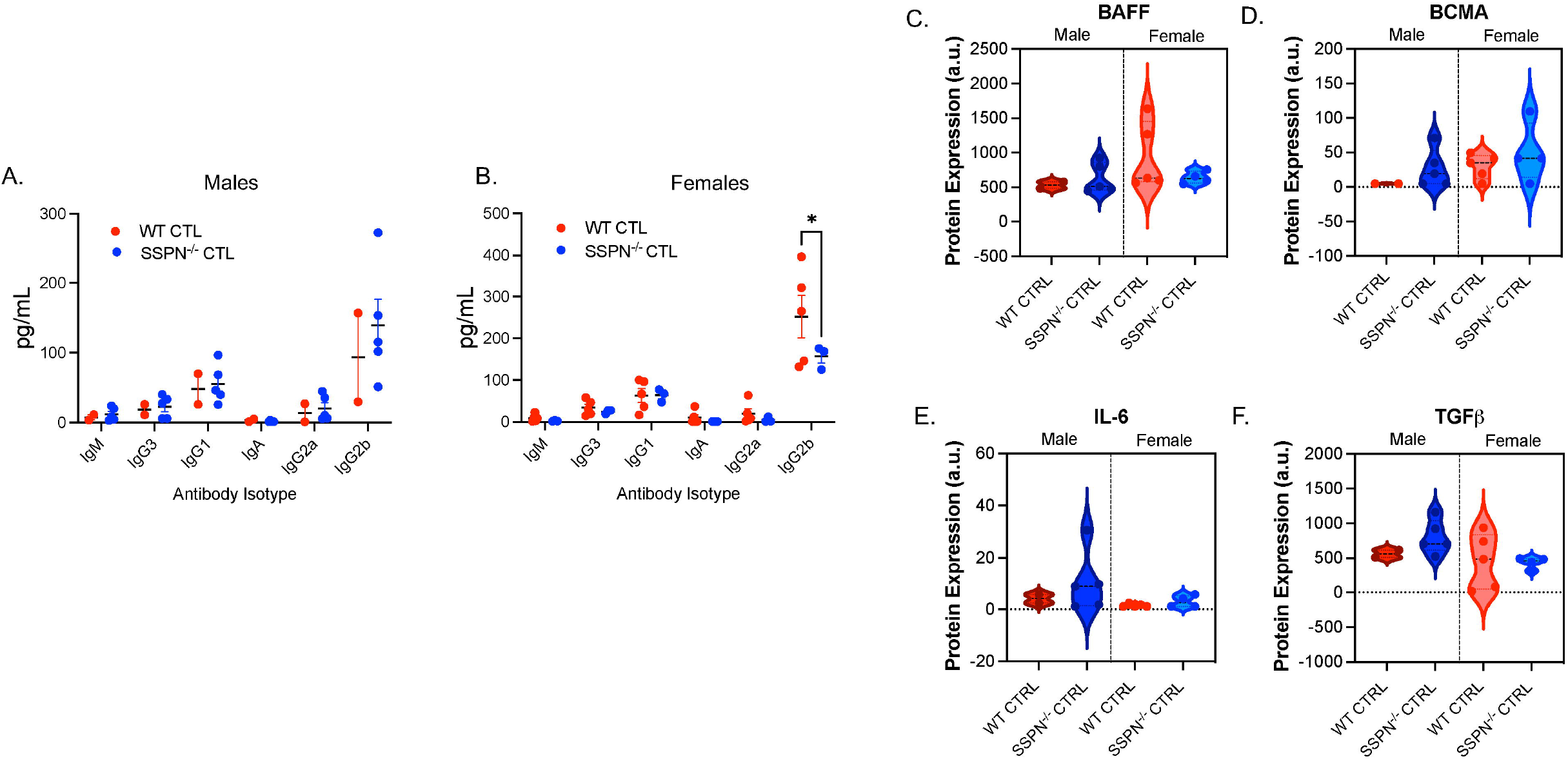
Assessment of Immunoglobulin Class Switching in Sarcospan-Deficient Mice. Blood serum was assessed for alterations in mouse immunoglobulin (Ig) subtypes (IgM, IgG3, IgG1, IgA, IgG2a and IgG2b) to determine whether essential B cell effector functions were affected by SSPN deletion in (A) male WT and SSPN^-/-^ blood serum and (B) female WT and SSPN^-/-^ blood serum. Blood serum was also assessed for important cytokines important for Ig switching (C) B cell activating factor (BAFF), (D) B cell maturation antigen (BCMA), (E) interleukin-6 (IL-6), and (F) transforming growth factor beta (TGFβ). Genotype differences in the male and female mouse groups were assessed separately using the two-tailed Student’s t-test and considered significant if *p* ≥ 0.05. Significant differences indicated by * for 0.05 and ** for 0.001.

The righthand panels in Figure 5C-F include serum cytokines important for B cell maturation and function to assess whether immunoglobulin switching in SSPN^-/-^ mice was affected by dysregulation of necessary factors that drive B cell maturation and isotype switching. It was found that serum levels of BAFF and BCMA are similar between groups shown in Figure 5C and D. These play crucial roles in B cell proliferation, survival, and activation. Whereas, in Figure 5E and 5F it shows that no major differences in serum levels of the pro-inflammatory IL-6 and pleiotropic TGF-β cytokine were found.

## 4. Discussion

Tetraspanin proteins are known for their ability to organize interacting protein partners within the cell membrane. These transmembrane proteins have emerged as having key roles in malignancy, the immune system and infectious disease [14]. Tetraspanins engage in numerous specific interactions that mediate formation of tetraspanin-enriched microdomains (TEMs) [14]. There are 33 tetraspanin proteins found in mammals [15]. Several tetraspanin proteins have been identified in leukocytes that may aid adhesion molecules in mediating immune cell recruitment and migration [8]. SSPN has conserved structures characteristic of tetraspanin proteins, with a large extracellular loop (LEL) that contains Cys residues that aid in maintenance of its tertiary structure [16]. However other tetraspanin features are lacking including several post-translational modification sites. The discovery of tetraspanin-like SSPN RNA message [13] and the presence of the protein in this study in several immune cell populations is intriguing. The protein interactions of SSPN within the DGC have been extensively characterized by Peter and colleagues in 2007 [17]. Subsequent studies showed that SSPN interacts with integrins and other proteins outside of the DGC [18,19]. SSPN+CD11b+ cells identified here could be of the myeloid lineage including monocytes/macrophages, neutrophils, eosinophils, and basophils but also include lymphoid lineage NK and peritoneal B-1 cells. Integrin αM (CD11b) is important for mediating adhesion of immune cells to endothelial cells and SSPN has been shown to important in muscle cell adhesion. Therefore, it could be that SSPN plays a role in adhesion in specific immune cells as well.

Hematopoiesis occurs in the bone marrow where stimulation of hematopoietic stems cells (HSC) produces multipotent progenitors that generate a range of lineage progenitor cells giving rise to the lymphoid and myeloid lineages [20]. The niche that hematopoietic stem cells reside in is replete with transcription factors, miRNAs and other mediators that provide an ideal microenvironment that supports their function. These factors balance HSC self-renewal and differentiation, activation and stemness. Using ImmuNet to examine functional relationships between SSPN and the immune system, it appears to have direct functional interactions with several hematopoiesis-linked factors with roles described below and marked in red in Figure 1B and numbered below.

SSPN was reported to have the strongest functional interaction with cathepsin O (CTSO). 1) CTSO is a cysteine cathepsin involved in cellular protein degradation with a role in modulating innate immune responsiveness and a less well characterized role in hematopoiesis [21]. Cathepsins are kept tightly in check to prevent uncontrolled proteolytic activity. Dysregulated cathepsin activity can contribute to cancer, neurodegeneration, and autoimmune diseases. 2) LRRC17 has defined roles in bone metabolism and regulation of osteoclast differentiation [22]. Interestingly, a human genome-wide association study linked variants in the *SSPN* gene with reduced bone density [23]. 3) ID4 belongs to the ID family of helix-loop-helix (HLH) proteins that lack the basic DNA binding domain [24]. ID proteins have been found necessary for normal differentiation and development of T cell, B cell, NK and innate lymphoid cells, dendritic cell, and myeloid cells. More recently ID protein have been found to be critical regulators of normal and leukemic hematopoietic stem and progenitor cells [24]. 4) PDLIM3 is a scaffolding protein that contains PDZ and LIM interaction domains that is involved in various signaling pathways including integrin-related, TGF-β, MAPK and NF-κB signaling pathways with a potential role in cancer development [25]. 5) ARHGEF6 is a Rho guanine nucleotide exchange factor which has been linked to acute myeloid leukemia (AML). AML is characterized by uncontrolled proliferation of myeloid progenitor cells leading to impaired myeloid differentiation and decreased percentage of normal blood cells [26]. Whereas, in hematopoiesis Rho GTPases are involved in regulating cell proliferation, differentiation, migration and self-renewal. 6) FBN1 is an extracellular matrix protein that normally restricts hematopoietic stem cell expansion however does not play a role in cell lineage commitment [27]. Fibrillin-1 also differentially modulates TGF-β activity in hematopoietic stem cell niches compared to erythroid niches [27]. 7) MFAP5 is a microfibril-associated glycoprotein that is a component of extracellular matrix myofibrils that promotes attachment of cells to microfibrils via alpha-V-beta-3-integrin [28] and deficiency of this gene causes neutropenia in mice. Future studies will need to be conducted to determine how SSPN interacts with these proteins and influences hematopoietic niches and the immune system.

The Human Protein Atlas data included in Figure 1E shows that *Sspn* RNA expression is altered as a function of B cell activation. In mice, B cells exist as three distinct subsets: B1, marginal zone (MZ), and B2 follicular (FO) [29]. Recent studies have shown that *Sspn* mRNA is expressed in B cells [29] [30]. Publicly available transcriptomics data indicate that *Sspn* transcript is increased as part of cluster 45 and is enriched in B1 B cells with diminished expression in marginal zone B cells (MZ B cells). The changes in SSPN expression in these distinct B cell subsets match the expression profile of the transcription factor *Myo1d* [30]. This suggests that SSPN may also be a MyoD regulated gene and supports earlier reports that it is a myoD target gene (ENCODE data set), see https://www.ncbi.nlm.nih.gov/gene/4654. MyoD is a nuclear protein in the basic helix-loop-helix transcription factor subfamily that regulates muscle cell differentiation and muscle regeneration. In addition, *Sspn* appears to be expressed as part of a gene cluster that is activated during B cell maturation and development [31]. Furthermore, *Sspn* expression in thymic B cells can be induced by AIRE, a long-noncoding RNA which also regulates expression of immune gene programs [32]. In our study we utilized extracellular flow cytometry and found that the SSPN protein is present in the membranes of B cells and CD11b+ and CD11c+ expressing cells (Figures 1C). In our study SSPN loss did not significantly alter B cell proliferation as shown in Figure 3D.

We investigated the impact of SSPN deficiency on immune cells of lymphoid lineage (Figure 3A and B) and Figure 4A, D, and G). Since currently the protein partners of SSPN in B cells and other immune cells are unknown it is interesting to note that in heart and skeletal muscle SSPN forms strong interactions with β1α7 integrins [4,18,19], dystroglycans, and sarcoglygans. B cells express different integrin isoforms that may affect integrin-mediated adhesion depending on their location including α4β, αLβ2, and αvβ3 [33] and changes in SSPN expression appear to accompany changes in integrin adhesion molecules. SSPN may play a variable role in modulating adhesion in B cells and other immune cells as well. B cells in this study identified as a CD19+B220+ population and were found to express SSPN. In mature B cells CD19 forms a multimolecular signal complex containing CD21, a complement receptor, and the tetraspanin molecule CD81. Our results suggest that SSPN may participate in signaling complexes that control other B cell functions or interactions [34]. In addition, there was a slight but significant increase %T cells in male SSPN^-/-^ splenocytes compared to WT, however it is unknown whether the quality of the T cell response or specific effector functions are affected including interferon-γ (IFN-γ) production, cytotoxic activity, proliferation, or induction of proliferation of other cells. There are no reports of *Sspn* transcript in NK cells and there were no significant differences in NK1.1^+^ expressing NK cells, however their expression trended lower in SSPN^-/-^ male and female mice. NK1.1 has been used to distinguish NK cells in specific murine strains, including C57BL/6 [35-37], [38]. The levels of an alternative NK cell type NKp46+ cells were also assessed to determine whether there were additional alterations in NK cell phenotypes in the SSPN^-/-^ mice.

Next, we investigated whether SSPN deficiency impacted myeloid lineage immune cells (Figure 4E and F). We examined plasmacytoid dendritic cells (pDCs), which are derived from myeloid progenitors that can be differentiated from DCs because of their rounded plasma cell shape that lacks projections. Both male and female SSPN^-/-^ splenocytes trended toward reduced numbers of pDCs. pDCs have DC-like functions and exist in lymphoid organs such as the spleen and are responsible for robust production of type I IFNs [39]. Therefore, they play important roles in antiviral immunity and have been implicated in initiation and progression of inflammation and inflammatory diseases. pDCs are continuously generated from hematopoietic stems cells through activation of both myeloid and lymphoid pathways [40]. It has been demonstrated that pDCs are critical for bridging innate and adaptive responses to systemic viral or bacterial infections. The slight reduction of pDCs in SSPN^-/-^ splenocytes may not have important consequences. Important future experiments to assess how SSPN deficiency impacts their pDC function will include pDC isolation and stimulation with influenza or CpG-2216 to assess cytokine secretion [41]. In contrast, DCs (CD11c+B220-) are significantly increased in SSPN^-/-^ female splenocytes. DC are the most efficient antigen presenting cells (APCs) in the immune system and have the unique ability to activate naïve T cells [42]. The spleen contains primarily non-migratory lymphoid tissue-resident DC that serve as APCs presenting incoming antigens to T cells [43]. Inappropriate responses to cell antigens can result from abnormal functionality of pDCs and DCs causing autoimmune diseases and chronic inflammatory conditions [44]. While increased DCs in SSPN^-/-^ female splenocytes does not signal abnormal function, it is an interesting observation and may warrant future investigation should SSPN^-/-^ mice exhibit signs of chronic immune system activation.

To further assess a role for SSPN in B cell function, we examined their important effector function - immunoglobulin isotype switching. Our study uncovered significantly lower baseline levels of the IgG2b antibody isotype in SSPN^-/-^ female mice. However, under immune challenge such as lipopolysaccharide (LPS) or LPS + IL-4, additional alterations in immune responses may be evident and differences in class switching recombination may be observable. Based on results in Figure 3B there were no alterations in B cell division therefore these differences are expected to result from division-independent mechanisms. To determine whether SSPN loss influenced the signals that promote B cell development we examined levels of B cell promoting factors including BAFF and BCMA (Figure 5C and D). The cytokine BAFF is a member of the TNF family and serves as a B cell survival factor that controls B cell maturation. Relatedly, a study examining BAFF-deficient mice found they lacked mature B cells [45]. Studies have shown that BAFF supports survival of several types of B cells, however not B1 B cells [45]. BCMA plays a critical role in regulating B cell proliferation, survival, and differentiation into plasma cells [46]. The IgSF surface glycoprotein CD19 is expressed on pro-B cells from the earliest B cell development stages until terminal differentiation as a plasma cell when its expression is lost. The presence of B cell promoting factors and our B cell differentiation results based upon CD19 expression suggest that SSPN does not regulate early B cell development and survival (Figure 3D). Human Atlas data in Figure 1E suggests that SSPN expression is significantly increased in switched memory B cells that memorize antigen characteristics that activated their parent B cell during initial infection to assure an accelerated, robust secondary immune response.

Other cytokines were examined with important roles in isotype switching including interleukin-6 (IL-6) and TGF-β. IL-6 is a proinflammatory cytokine produced by APCs and nonhematopoietic cells. In vitro studies indicated that IL-6 is a B cell growth factor that induces plasma cell differentiation [47]. Subsequent in vivo studies showed it has an important role in class switching and antibody production [47]. TGF-β is a pleiotropic cytokine involved in both suppressive and inflammatory immune responses, which has been shown to be important in B cell IgA isotype expression when stimulated with lipopolysaccharide (LPS), a bacterial cell wall component. Under baseline conditions TGF-β provides only partial or incomplete IgA switching signals keeping levels of these isotypes low. A limitation of our study is that we are examining the impact of SSPN deletion on baseline isotype abundance in the absence of immune stimulus. Future studies will be directed towards examining the role of SSPN in antibody class switching in response to different immune stimuli.

## 5. Conclusions

Our study is the first to examine a role for SSPN in immune cell production and function and explore the functional link between SSPN and factors that regulate bone marrow cell niche, hematopoietic lineages, and the immune system response. We showed that SSPN is expressed on the surface of B cells, CD11b+, and CD11c+ cells. Loss of SSPN did not disrupt the production of the immune cell populations assessed in this study including B cells, T cells, pDCs, DCs, and NK cells. However, a few alterations in immune cell abundance were noted in SSPN^-/-^ mice including higher % T cells in male and increased % pDCs in female splenocytes. Immunoglobulin class switching was assessed in non-stimulated B cells and the levels of isotype IgG2b were significantly lower in SSPN^-/-^ compared to WT female mice. Overall, these findings suggest that SSPN plays distinct yet unknown roles in immune cell and hematopoietic cell functions.

## CRediT Author Contribution Statement

**Isela Valera:** - Data curation. Writing – review and editing. **Rhiannon Crawford:** Data curation. **Kyle Smith:** - Data curation **Aida Rahimi Kahmini:** Data curation. **Michelle Parvatiyar:** – Writing – original draft, review & editing, Project administration, Funding acquisition.

## Declaration of Competing Interest

We declare that the authors have no conflict of interest concerning this paper.

## Acknowledgements

The authors thank the AHA Scientist Development Grant 16SDG29120002 for support while developing this project. The authors have no conflicts to disclose. The content here within is solely the author’s responsibility and does not necessarily represent the official views of the Florida State University.

## Notes

### Competing Interest Statement

The authors have declared no competing interest.

